# Prolonged low-dose dioxin exposure impairs metabolic adaptability to high-fat diet feeding in female but not male mice

**DOI:** 10.1101/2020.09.12.294587

**Authors:** Geronimo Matteo, Myriam P Hoyeck, Hannah L Blair, Julia Zebarth, Kayleigh RC Rick, Andrew Williams, Rémi Gagné, Julie K Buick, Carole L Yauk, Jennifer E Bruin

## Abstract

**Objective:** Human studies consistently show an association between exposure to persistent organic pollutants, including 2,3,7,8-tetrachlorodibenzo-*p*-dioxin (TCDD, aka “dioxin”), and increased diabetes risk, but rarely consider potential sex differences. We previously showed that a single high-dose TCDD exposure (20 µg/kg) decreased plasma insulin levels in both male and female mice *in vivo*, but effects on glucose homeostasis were sex-dependent. The purpose of the current study was to determine whether prolonged exposure to a physiologically relevant low-dose of TCDD impacts glucose homeostasis and/or the islet phenotype in a sex-dependent manner in either chow-fed or high fat diet (HFD)-fed mice.

**Methods:** Male and female mice were exposed to 20 ng/kg/d TCDD 2x/week for 12 weeks and simultaneously fed standard chow or a 45% HFD. Glucose homeostasis was assessed by glucose and insulin tolerance tests, and glucose-induced plasma insulin levels were measured *in vivo*. Histological analysis was performed on pancreas from male and female mice, and islets were isolated from females at 12 weeks for Tempo-Seq® analysis.

**Results:** Low-dose TCDD exposure did not lead to adverse metabolic consequences in chow-fed male or female mice, or in HFD-fed males. However, TCDD accelerated the onset of HFD-induced hyperglycemia and impaired glucose-induced plasma insulin levels in female mice. TCDD caused a modest increase in islet area in males irrespective of diet, but reduced % beta cell area within islets in females. RNAseq analysis of female islets also revealed abnormal changes to endocrine and metabolic pathways in TCDD-exposed HFD-fed females compared chow-fed females.

**Conclusions:** Our data suggest that prolonged low-dose TCDD exposure has minimal effects on glucose homeostasis and islet morphology in chow-fed male and female mice, but promotes maladaptive metabolic responses in HFD-fed females.

## 1. Introduction

Global diabetes incidence is on the rise, yet the underlying causes for this increase remain to be elucidated (1). Type 2 diabetes, the most common form of diabetes, is characterized by chronic hyperglycemia, insufficient insulin production by pancreatic beta cells, and peripheral insulin resistance (2). Genetics and lifestyle factors, such as physical inactivity and poor diet, are known risk factors for type 2 diabetes (2), but cannot alone account for the rapid increase in diabetes burden.

Persistent organic pollutants (POPs) are particularly concerning environmental contaminants since they resist degradation and bioaccumulate. High serum POP concentrations are positively associated with a modest relative risk of developing type 2 diabetes (3–14) and insulin resistance (15,16) both in the general population with chronic low-dose POP exposure (11–13) and in individuals with high-dose exposure (e.g. veterans, chemical disaster victims, and occupational workers) (5,6,14). Most epidemiological studies that investigate the association between POP exposure and diabetes risk only report male data. Interestingly, the few studies that compared sexes suggest a stronger link between POP serum concentrations and diabetes risk in females than males. For example, studies on the Seveso population exposed to 2,3,7,8-tetrachlorodibenzo-*p*-dioxin (TCDD, aka “dioxin”) reported a higher relative risk of developing diabetes in females than males (5,17). In addition, high serum POP levels in the Michigan (18) and Yucheng (10) cohorts were associated with increased diabetes incidence in women only. A causal link between POP exposure and diabetes pathogenesis remains to be established, and potential sex differences require further investigation.

Dioxin/TCDD is a potent aryl hydrocarbon receptor (AhR) agonist and an excellent model chemical for investigating the relationship between dioxin-like environmental contaminants and diabetes risk. Although TCDD is now well regulated worldwide, there are hundreds of dioxin-like chemicals that remain a global concern for human health (19). We have shown that TCDD exposure induces AhR-regulated genes, including cytochrome P450 1a1 (*Cyp1a1*), in pancreatic islets, demonstrating that TCDD reaches the endocrine pancreas *in vivo* (20). Direct TCDD exposure also reduced glucose-stimulated insulin secretion in mouse and human islets *in vitro* (20). Interestingly, a single high-dose injection of TCDD (20 µg/kg) *in vivo* reduced plasma insulin levels in both male and female mice, but effects on overall glucose homeostasis differed drastically between sexes (21). TCDD-exposed males had modest fasting hypoglycemia for ~4 weeks post-injection, increased insulin sensitivity, and decreased % beta cell area within islets compared to vehicle-exposed controls. Conversely, TCDD did not alter insulin sensitivity or islet composition in females, but induced transient hyperglycemia during a glucose tolerance test (GTT) at 4 weeks post-injection (21). Whether similar sex-specific effects occur with a physiologically relevant dose of TCDD remains to be investigated. Interestingly, we found that transient low-dose TCDD exposure (20 ng/kg/d, 2x/week) during pregnancy/lactation in female mice did not impact glucose homeostasis in chow-fed females, but promoted high fat diet (HFD)-induced obesity and hyperglycemia post-exposure (22). It remains unknown whether prolonged low-dose TCDD exposure has a similar interaction with HFD feeding in non-pregnant female mice, and whether the susceptibility to hyperglycemia is sex-specific. The purpose of the current study was to determine whether prolonged exposure to a physiologically relevant low-dose of TCDD impairs glucose homeostasis and/or the islet phenotype in a sex-dependent manner in either chow-fed or HFD-fed mice.

## 2. Material and Methods

### 2.1 Animals

Male and female C57BL/6 mice, 6-8 weeks old (Charles River; Raleigh, NC, USA), were maintained on a 12-hour light/dark cycle. All mice received *ad libitum* access to a standard chow diet for 1 week prior to starting experimental treatments. All experiments were approved by Carleton University and University of Ottawa Animal Care Committees and carried out in accordance with Canadian Council on Animal Care guidelines. All experimental groups were matched for mean body weight and fasting blood glucose levels prior to starting treatments to ensure that no group was statistically different from another.

Male and female mice received intraperitoneal (i.p.) injections of corn oil (CO; vehicle control, 25 ml/kg) (#C8267-2.5L, Sigma Aldrich, St. Louis, MO, USA) or a low-dose of TCDD (20 ng/kg/d) (Sigma Aldrich, # 48599) 2x/week for 12 weeks (**Fig. 1A**). Simultaneously, mice were fed either standard rodent chow (20% fat, 66% carbohydrate, 14% protein; #2918, Harlan Laboratories, Madison, WI, USA) or a HFD (45% fat, 35% carbohydrate, 20% protein; #D12451, Cedarlane, Burlington, ON, Canada), generating the following experimental groups (n=10 per group per sex): COChow, COHFD, TCDDChow, TCDDHFD. The TCDD dosing protocol was selected based on studies showing that a similar dosing regimen induced *Cyp1a1* mRNA and enzyme activity in liver/lung tissues, and produced an environmentally relevant steady-state tissue burden without overt TCDD toxicity (e.g. weight loss, immunosuppression, hepatic toxicity) (23,24). More specifically, circulating TCDD concentrations were consistently ~10 pg/g after 2 years of chronic TCDD (22 ng/kg/d) exposure (23), and ~8.8 pg/g after 15 weeks of prolonged TCDD (46 ng/kg/d) exposure in rats (24); these circulating TCDD levels are within the range of background dioxin levels reported in the United States (≤ 10 pg/g) and corresponds to individuals in the upper quartile (≥ 5.2 pg/g) with increased diabetes prevalence (25). Rats provide a useful model for dose selection since TCDD biodistribution is similar between mice and rats (24,26,27). We have previously shown that this TCDD dosing protocol significantly induced *Cyp1a1* in mouse islets and liver after 2 weeks (20) and was generally well tolerated in mice (e.g. no weight loss) (22). There was no difference in the degree of *Cyp1a1* induction in islets by i.p. administration versus oral gavage (data not shown). Therefore, i.p. injection was used rather than oral administration since it allowed us to control the spread of the chemical in animal cages and the dose of TCDD consumed by each mouse.

**Fig. 1.**
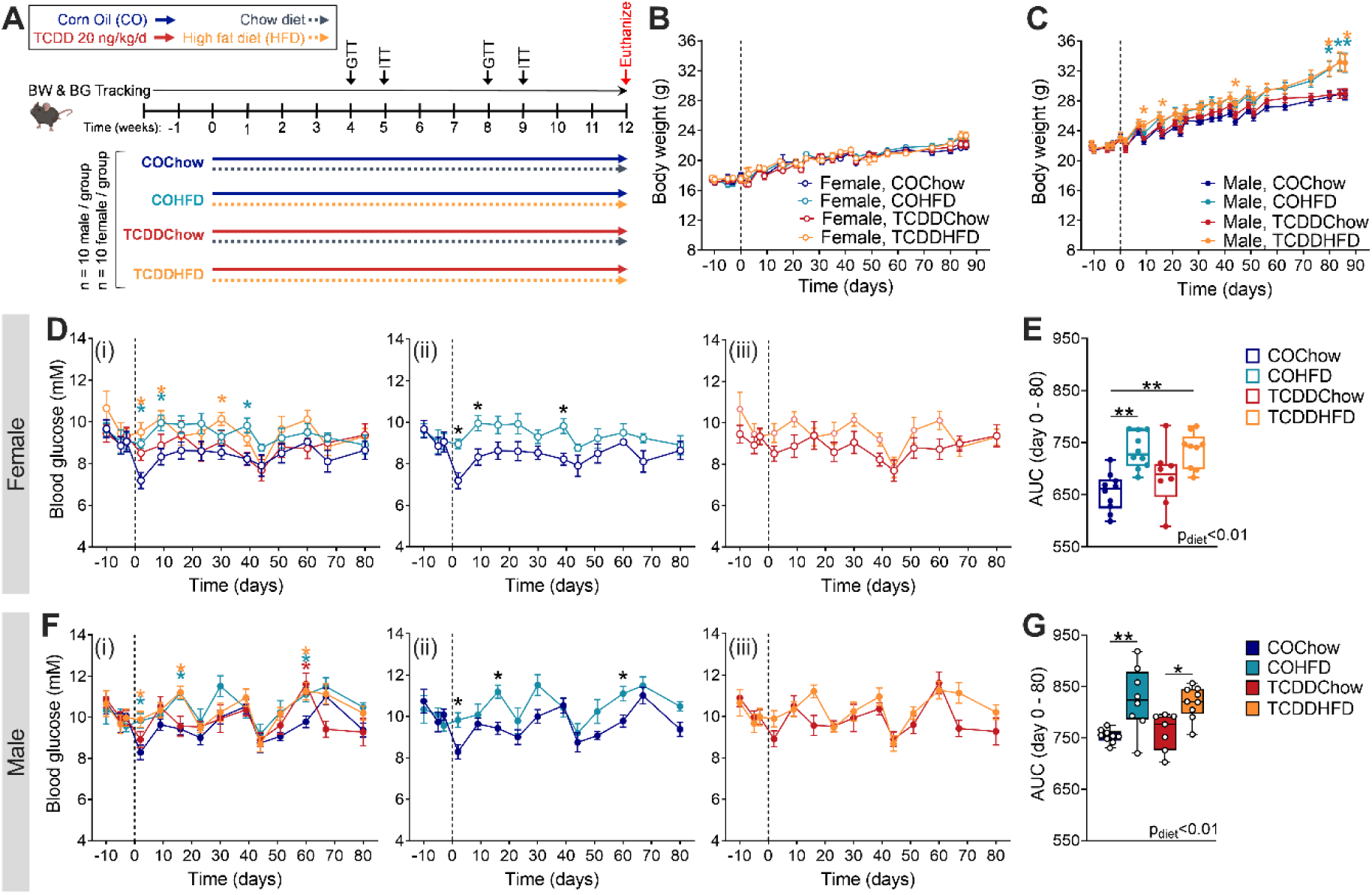
Low-dose TCDD exposure does not impact body weight or fasting blood glucose levels in mice. (**A**) Male and female mice were exposed to corn oil or 20 ng/kg/d TCDD 2x/week for 12 weeks, and simultaneously fed either a chow diet or 45% HFD. Body weight and blood glucose was tracked throughout the 12-week study. BW = body weight; BG = blood glucose; GTT = glucose tolerance test; ITT = insulin tolerance test. (**B-C**) Body weight and (**D-G**) blood glucose were measured weekly following a 4-hour morning fast in (**B, D, E**) females and (**C, F, G**) males (n=7-10 per group). (**D, F**) Blood glucose data are presented as (**i**) all groups compared to COChow, (**ii**) COHFD versus COChow, and (**iii**) TCDDHFD compared to TCDDChow. Data are presented as mean ± SEM in line graphs or min/max values in box and whisker plots. Individual data points on box and whisker plots represent biological replicates (different mice). *p <0.05, **p <0.01, coloured stars are versus COChow. The following statistical tests were used: (**B, C, D, F**) two-way REML-ANOVA with Tukey’s multiple comparisons test; (**E, G**) two-way ANOVA with Tukey’s multiple comparisons test.

Whole pancreas was harvested from a subset of male and female mice at week 12 of the study and stored in 4% paraformaldehyde (PFA; #AAJ19943K2, Thermo Fisher Scientific, Waltham, MA, USA) for 24-hours, followed by long-term storage in 70% ethanol for histological analysis (n=4-5 per group per sex). Pancreatic islets were isolated from a different subset of female mice at week 12 and stored in RNAlater (#76106, Qiagen, Hilden, Germany) for analysis by quantitative real-time PCR (qPCR) and TempO-Seq® (n = 3-5 per group).

### 2.2 Metabolic Assessments

All metabolic assessments were performed on conscious, restrained mice. Blood samples were collected via the saphenous vein using heparinized microhematocrit tubes. Blood glucose was measured using a handheld glucometer (Johnson & Johnson, New Brunswick, NJ, USA).

Body weight and blood glucose measurements were performed weekly following a 4-hour morning fast (approximately 8am to 12pm). For all metabolic assessments, time 0 indicates the blood collected prior to glucose or insulin administration. For GTTs, mice received an i.p. bolus of glucose (2 g/kg) following a 4-hour morning fast. Blood samples were collected at 0, 15, and 30 minutes for measuring plasma insulin levels by ELISA (#80-INSMSU-E01, RRID:AB_2792981, ALPCO, Salem, NH, USA). For insulin tolerance tests (ITT), mice received an i.p. bolus of insulin (0.7 IU/kg, #02024233, Novo Nordisk, Toronto, Canada) following a 4-hour morning fast.

### 2.3 Immunofluorescence Staining and Image Quantification

Tissues were processed and embedded in paraffin wax blocks (University of Ottawa Heart Institute, Ottawa, ON, Canada) and then sectioned (5 µm). Immunofluorescence staining was performed as previously described (20). In brief, slides were deparaffinized with sequential incubations in xylene and ethanol. Heat-induced epitope retrieval was performed in 10 mM citrate buffer at 95°C for 10-15 minutes, and slides were incubated with Dako Serum Free Protein Block (#X090930-2, Agilent, Santa Clara, CA, USA) at room temperature. Slides were incubated overnight at 4°C with primary antibodies, and then for 1-hour at room temperature with secondary antibodies. Coverslips were mounted with Vectashield® hardset mounting medium with DAPI (#H-1500, RRID:AB2336788, Vector Laboratories, Burlingame, CA, USA).

The following primary antibodies were used: rabbit anti-insulin (1:200, C27C9, #3014, RRID:AB_2126503, Cell Signaling Technology), and mouse anti-glucagon (1:1000; #G2654, RRID:AB_259852, Sigma Aldrich). The following secondary antibodies were used: goat anti-mouse IgG (H+L) secondary antibody, Alexa Fluor 488 (1:1000, #A11029, RRID:AB_138404, Invitrogen); and goat anti-rabbit IgG (H+L) secondary antibody, Alexa Fluor 594 (1:1000, #A11037, RRID:AB_2534095, Invitrogen).

Whole pancreas sections were imaged using a Zeiss Axio Observer 7 microscope, and immunofluorescence was manually quantified using Zen Blue 2.6 software (Zeiss, Oberkochen, Germany). For all measurements, a range of 4-36 islets per mouse were quantified and the average was reported for each biological replicate. The % hormone^+^ area per islet was calculated as [(hormone^+^ area / islet area) x 100].

### 2.4 Islet Isolation

Islets were isolated by pancreatic duct injections with collagenase (1000 U/ml; #C7657, Sigma-Aldrich), as previously described (20). In brief, inflated pancreas tissue was excised and incubated at 37°C for 12 minutes, and the collagenase reaction quenched with cold Hank’s balanced salt solution (HBSS) with 1 mM CaCl_2_. Pancreas tissue was washed in HBSS+CaCl_2_, resuspended in Ham’s F-10 (#SH30025.01, HyClone, GE Healthcare Biosciences, Pittsburgh, PA, USA) containing 7.5% BSA (#10775835001, Sigma-Aldrich) and 1% Penicillin-Streptomycin (#30-002-CI, Corning, Tewksbury, MA, USA), and filtered through a 70 μm cell strainer (#07-201-431, Corning). Islets were handpicked under a dissecting scope to >95% purity.

### 2.5 Quantitative Real Time PCR

RNA was isolated from pancreatic islets using the RNeasy Micro Kit (#74004, Qiagen), as per the manufacturer’s instructions, with the following amendment: 7 volumes of buffer RLT + DTT were added to the samples prior to lysing with 70% EtOH. RNA quality was assessed using a Nanodrop®. DNase treatment was performed prior to cDNA synthesis with the iScript™ gDNA Clear cDNA Synthesis Kit (#1725035, Bio-Rad, Mississauga, ON, Canada). qPCR was performed using SsoAdvanced Universal SYBR Green Supermix (Bio-Rad, #1725271) and run on a CFX96 or CFX394 (Bio-Rad). *Ppia* was used as the reference gene since it displayed stable expression under control and treatment conditions. Data were analyzed using the 2^−ΔΔCT^ relative quantitation method. Primer sequences are listed in **Supp. Table 1** (28).

### 2.6 TempO-Seq®

Gene expression was measured in female islets (n = 3-4 per group) at week 12 of the study using the TempO-Seq® Mouse Whole Transcriptome panel in a 96-well format, according to the TempO-Seq® Assay User Guide (BioSpyder Technologies Inc, Carlsbad, CA). Briefly, 2 µl of whole islet RNA was diluted with 2 µl of 2x TempO-Seq lysis buffer, resulting in samples ready for assay input in 1x TempO-Seq® Enhanced Lysis Buffer (RNA input range = 15.5 - 98 ng/µl, all within recommended limits for whole transcriptome TempO-Seq® assays). Experimental samples, in addition to a positive control (100 ng/µl of Universal Mouse Reference RNA; Agilent, cat # 750600) and a negative control (1x TempO-Seq® Enhanced Lysis Buffer alone), were hybridized to the Detector Oligo (DO) Pool in an Annealing Mix for panels with > 5,000 probes for 10 minutes at 70°C followed by a temperature gradient with a ramp rate of 0.5°C/minute to 45°C over 50 minutes with a 16-hour hold at 45°C and then cooled to 25°C. Hybridization was followed by nuclease digestion to eliminate excess, unbound, or incorrectly bound DOs at 37°C for 90 minutes. A pool of amplification templates was then generated by ligating the DO pairs bound to adjacent target sequences for 60 minutes at 37°C, followed by a 15-minute enzyme denaturation at 80°C. The control and experimental amplification templates (10 µl of ligated DOs) were pipetted into their respective wells of the 96-well PCR Pre-Mix and Primer plate. Amplification proceeded using a CFX96 Real-Time PCR Detection System (Bio-Rad) to incorporate a unique sample index/tag sequence and the sequencing adaptors for each sample using the following PCR settings: 37°C for 10 minutes, 95°C for 1 minute; 25 cycles of 95°C for 10 seconds, 65°C for 30 seconds, 68°C for 30 seconds (with optical read for visual sample QC); 68°C for 2 minutes; hold at 25°C prior to library pooling and purification. Library purification was achieved by pooling and purifying labelled amplicons using the NucleoSpin® Gel and PCR Clean-up kits (Takara Bio USA, Inc, Mountain View, CA USA) with three modifications to the NucleoSpin® User Manual, including the addition of a second wash with buffer NT3, an increase of the spin-to-dry step from 1 minute to 10 minutes, and an additional elution with NE buffer to maximize the library yield. The pooled and purified library was quantified using the Universal KAPA Library Quantification kit for Illumina platforms (Roche, # KK4824). The pooled TempO-Seq® library was sequenced in-house using the NextSeq® 500/550 High Output Kits v2 (75 cycles) on an Illumina NextSeq 500 High-Throughput Sequencing System (Illumina, San Diego, CA, USA). A median read depth of 2 million reads/sample was achieved.

Reads were extracted from the BCL files, demultiplexed (i.e. assigned to respective sample files) and were processed into FASTQ files with bcl2fastq v.2.17.1.14. The FASTQ files were processed with the “pete.star.script_v3.0” supplied by BioSpyder. Briefly, the script uses star v.2.5 to align the reads and the qCount function from QuasR to extract the feature counts specified in a GTF file from the aligned reads. The data were then passed through internal quality control scripts. Boxplots of the log2 CPM (counts per million) were plotted to ensure a reproducible distribution between replicates within a group. Hierarchical clustering plots were generated (hclust function: default linkage function of hclust function in R; complete-linkage) using a distance metric defined as 1-Spearman correlation in order to identify potential outliers. Probes with low counts (i.e. less than a median of 5 counts in at least one group) were flagged as absent genes and eliminated from the dataset. Differentially expressed gene (DEG) analysis was conducted using the R software (29) on the counts using the default parameters of DESeq2 (30) with respective control and exposure groups. A shrinkage estimator was applied to the fold change estimates using the apeglm method (31) using the lfcShrink() function.

Probes reaching the threshold of an adjusted p-value < 0.05 and an absolute fold change > 1.5 (on a linear scale) were defined as DEGs and were retained for pathway analysis. Gene set analysis using KEGG pathways was conducted using DAVID (32,33). Pathways with a modified Fisher’s exact p-value < 0.01 were considered enriched. Data are presented as bar graphs showing pathway fold enrichment, and secondly, as bar graphs with DEG counts for pathways that were trending (p = 0.01 - 0.1) or statistically enriched (p < 0.01). Pathways that were not trending or statistically enriched are indicated on bar graphs as not having the DEG count minimum threshold for significance.

### 2.7 Statistical Analysis

Statistical analysis was performed using GraphPad Prism Software (GraphPad Software Inc., La Jolla, CA, RRID:SCR_002798). Specific statistical tests are indicated in the figure legends and p < 0.05 was considered statistically significant for all analyses. Statistically significant outliers were detected by a Grubbs’ test with α = 0.05. All data were tested for normality using a Shapiro-Wilk test and for equal variance using either a Brown-Forsyth test (for one-way ANOVAs) or an F test (for unpaired t tests). Non-parametric statistics were used in cases where the data failed normality or equal variance tests. Parametric tests were used for all two-way ANOVAs, but normality and equal variance were tested on area under the curve values and by one-way ANOVAs. Data in line graphs are presented as mean ± SEM. Data in box and whisker plots are displayed as median, with whiskers representing maximum and minimum values. Each individual data point represents a different biological replicate (i.e. individual mouse).

### 2.8 Data and Resource Availability

The datasets generated and/or analyzed during the current study are available from the corresponding author upon reasonable request. TempO-Seq® data have been submitted to National Centre for Biotechnology Information (NCBI) Gene Expression Omnibus (GEO) and are accessible under accession number GSE144765.

## 3. Results

### 3.1 Low-dose TCDD exposure does not impact body weight or fasting blood glucose levels in mice

Male and female mice were exposed to CO (vehicle) or 20 ng/kg/d TCDD 2x/week for 12 weeks, and simultaneously fed either a chow diet or 45% HFD (**Fig. 1A**). We were particularly interested in whether low-dose TCDD exposure alone would promote weight gain (i.e. COChow versus TCDDChow) and/or accelerate diet-induced obesity (i.e. COHFD versus TCDDHFD), as previously seen in female mice exposed to TCDD during pregnancy (22). Interestingly, TCDD exposure had no effect on body weight in chow-fed females in this study (**Fig. 1B**). Female mice were also resistant to HFD-induced weight gain within the 12-week study timeframe, regardless of chemical exposure (**Fig. 1B**). Likewise, TCDD exposure did not alter body weight in chow-fed or HFD-fed male mice, but HFD feeding promoted significant weight gain in male mice after 80 days (**Fig. 1C**). TCDD did not influence fasting blood glucose levels in any group, but HFD feeding caused fasting hyperglycemia in both CO- and TCDD-exposed females (**Fig. 1D,E**) and males (**Fig. 1F,G**).

### 3.2 TCDD exposure accelerates the onset of HFD-induced glucose intolerance in female mice

TCDD had no effect on glucose tolerance in chow-fed female or male mice at weeks 4 or 8 of the study (COChow versus TCDDChow; **Fig. 2**). However, we observed an interaction between TCDD exposure and HFD feeding in female mice (**Fig. 2A-D**). At week 4, HFD feeding had no impact on glucose tolerance in CO-exposed females (COChow versus COHFD; **Fig. 2Aii,2B**), but TCDDHFD females were significantly hyperglycemic at 15- and 30-minutes post-glucose bolus and had a significantly elevated overall glucose excursion compared to TCDDChow females (**Fig. 2Aiii,2B**). By week 8, both COHFD and TCDDHFD females were glucose intolerant compared to chow-fed females (**Fig. 2C,D**). In contrast, HFD-fed male mice were significantly hyperglycemic compared to chow-fed males during the GTTs at weeks 4 and 8, irrespective of chemical exposure (**Fig. 2E-H**). In other words, TCDD-exposed and CO-exposed males showed similar responses to HFD feeding (**Fig. 2E-H**), whereas HFD-fed female mice had accelerated onset of hyperglycemia in the presence of TCDD (**Fig. 2A-D**).

### 3.3 TCDD-exposed female mice lack a compensatory hyperinsulinemic response to HFD feeding but show normal insulin tolerance in vivo

There were no changes in fasting or glucose-induced plasma insulin levels in TCDD-exposed or HFD-fed female and male mice at week 4 of the study (**Supp. Fig. 1A-B**), but at week 8 TCDD-exposed females showed an impaired insulin response to HFD feeding compared to CO-exposed females. Specifically, COHFD females showed an expected compensatory increase in plasma insulin levels during the GTT at week 8 (**Fig. 3A**), with ~2.3-fold more insulin overall than COChow females (**Fig. 3A-AUC**). In contrast, plasma insulin levels in TCDDHFD females were comparable to COChow and TCDDChow females (**Fig. 3A**). TCDD did not impact insulin levels in either chow-fed or HFD-fed male mice at week 8 (**Fig. 3B**), but rather diet had an overall effect on plasma insulin levels in males (**Fig. 3B**). These results suggest that TCDD exposure prevents HFD-induced hyperinsulinemia in females only.

**Fig. 2.**
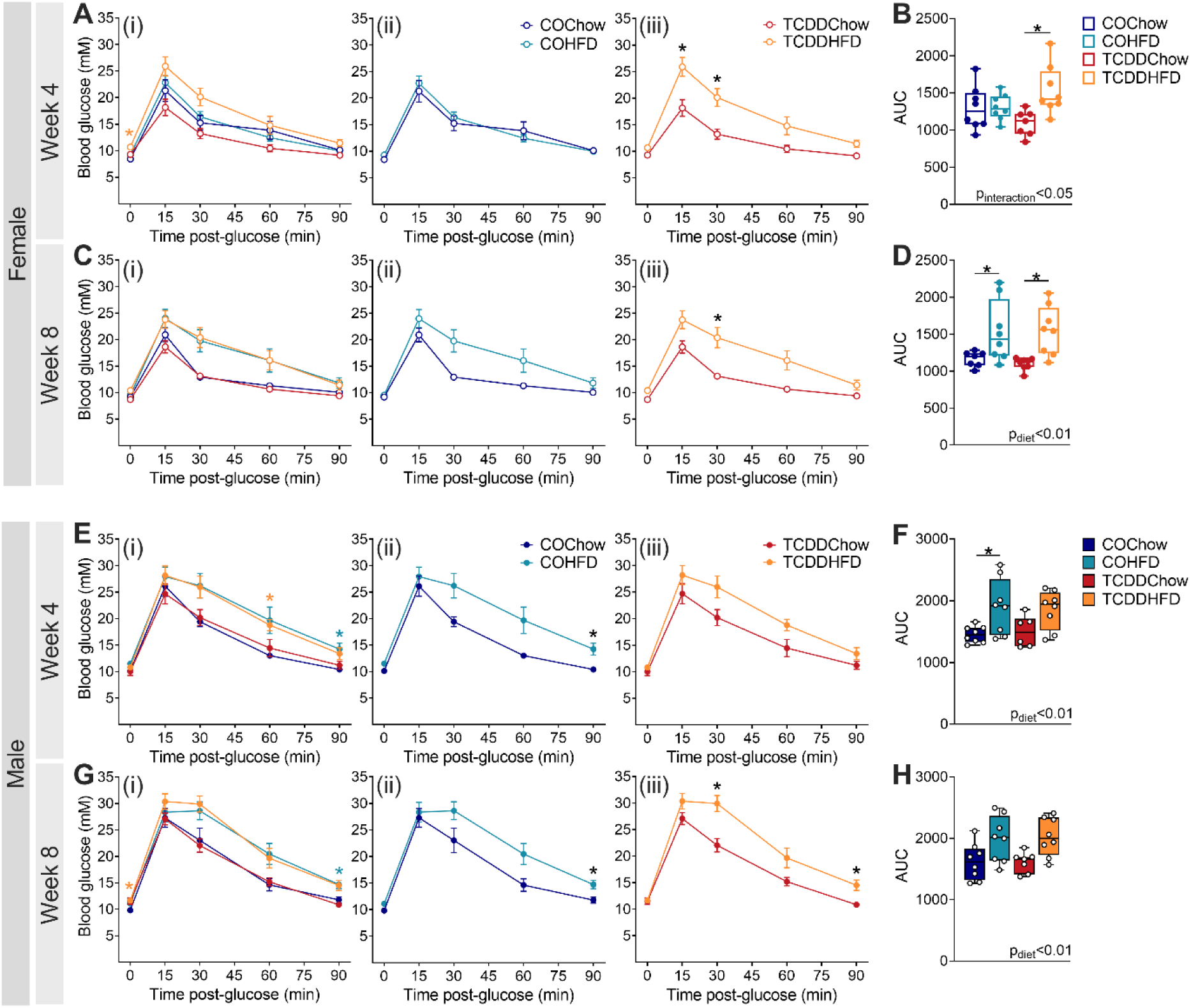
TCDD exposure accelerates the onset of HFD-induced glucose intolerance in female mice. Glucose tolerance tests were performed after 4 and 8 weeks of TCDD exposure with/without HFD feeding (see **Fig. 1A** for study timeline). Blood glucose levels in (**A-D**) females and (**E-H**) males at (**A, B, E, F**) week 4 and (**C, D, G, H**) week 8 (n=6-8 per group). (**A, C, E, G**) Blood glucose data are presented as (**i**) all groups compared to COChow, (**ii**) COHFD versus COChow, and (**iii**) TCDDHFD compared to TCDDChow. Data are presented as mean ± SEM in line graphs or min/max values in box and whisker plots. Individual data points on box and whisker plots represent biological replicates (different mice). *p <0.05, coloured stars are versus COChow. The following statistical tests were used: (**A, C, E, G**) two-way RM ANOVA with Tukey’s multiple comparison test; (**B, D, F, H**) two-way ANOVA with Tukey’s multiple comparison test.

**Fig. 3.**
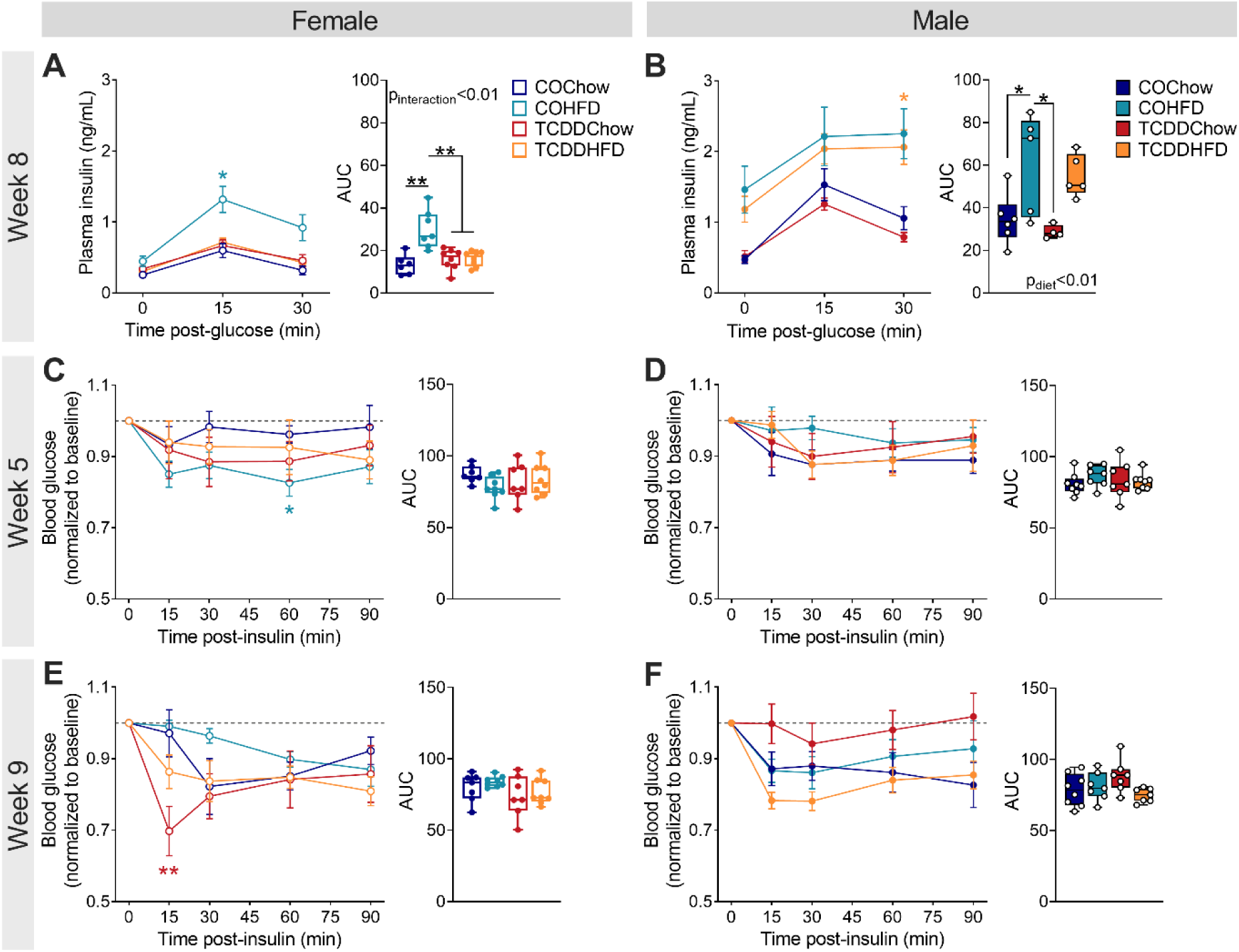
TCDD-exposed female mice lack an appropriate compensatory hyperinsulinemic response to HFD feeding but show normal insulin tolerance. Glucose-stimulated plasma insulin levels were assessed *in vivo* at week 8, and insulin tolerance at week 5 and 9 (see **Fig. 1A** for study timeline). Plasma insulin levels during a GTT at week 8 of the study in (**A**) females and (**B**) males (n=5-8 per group). Blood glucose levels during an ITT at (**C, D**) 5 weeks and (**E, F**) 9 weeks in (**C, E**) females and (**D, F**) males (n=7-8 per group). Data are presented as mean ± SEM in line graphs or min/max values in box and whisker plots. Individual data points on box and whisker plots represent biological replicates (different mice). *p <0.05, **p <0.01, coloured stars are versus COChow. The following statistical tests were used: (**A-F**) line graphs, two-way RM ANOVA with Tukey’s multiple comparison test; box and whisker plots, two-way ANOVA with Tukey’s multiple comparison test.

To determine whether changes in peripheral insulin sensitivity were also contributing to the accelerated hyperglycemia in TCDDHFD females, we performed ITTs at weeks 5 and 9. HFD-fed female and male mice were not overtly insulin resistant in this study (**Fig. 3C-F**). At week 5, COHFD females were slightly more insulin sensitive than COChow females at 60 minutes after the insulin bolus (**Fig. 3C**) and at week 9 TCDDChow females were modestly more insulin sensitive at 15 minutes post-bolus compared to COChow females (**Fig. 3E**), but there were no changes in overall blood glucose levels during the ITTs (**Fig. 3C-F AUC**).

We measured expression of genes involved in insulin-dependent lipogenesis and gluconeogenesis pathways in the liver at week 12 as another indicator of insulin sensitivity. In females, HFD feeding significantly downregulated genes involved in gluconeogenesis (**Supp. Fig. 2A**) (28), including *G6pc* (glucose-6-phosphatase catalytic subunit), *Ppargc1a* (peroxisome proliferator-activated receptor gamma coactivator 1 alpha), and *Pck1* (phosphoenolpyruvate carboxykinase 1). There was a significant interaction between TCDD and HFD exposure on expression of *Acacb* (acetyl-CoA carboxylase beta; **Supp. Fig. 2A**) (28), but this was modest and unlikely to explain the differences in plasma insulin levels or glucose homeostasis in female mice. There were no differences in any of the measured genes in the liver of male mice following TCDD and/or HFD exposure (**Supp. Fig. 2B**) (28).

### 3.4 TCDD exposure has sex-specific effects on islet histology in mice

To investigate why TCDD-exposed female mice did not develop hyperinsulinemia following HFD feeding, we examined islet size and endocrine cell composition in the pancreas at week 12. We predicted that HFD-fed mice might show increased islet size compared to chow-fed mice, but this was not the case in either sex (**Fig. 4A,D,G**). Instead, there was an unexpected significant overall effect of TCDD exposure to increase average islet size in male mice (**Fig. 4D**, p_chemical_ < 0.05), but not females (**Fig. 4A**). On the other hand, TCDD exposure significantly reduced the percentage of islet area immunoreactive for insulin in females (**Fig. 4B,G**), but not males (**Fig. 4E,G**). The effect of TCDD on islet composition in females was independent of diet and therefore does not explain the interactive effect between TCDD and HFD on plasma insulin levels in females. TCDD and/or HFD exposure had no effect on insulin^+^ area in male islets (**Fig. 4E,G**) or on glucagon^+^ area in either sex (**Fig. 4C,F,G**).

**Fig. 4.**
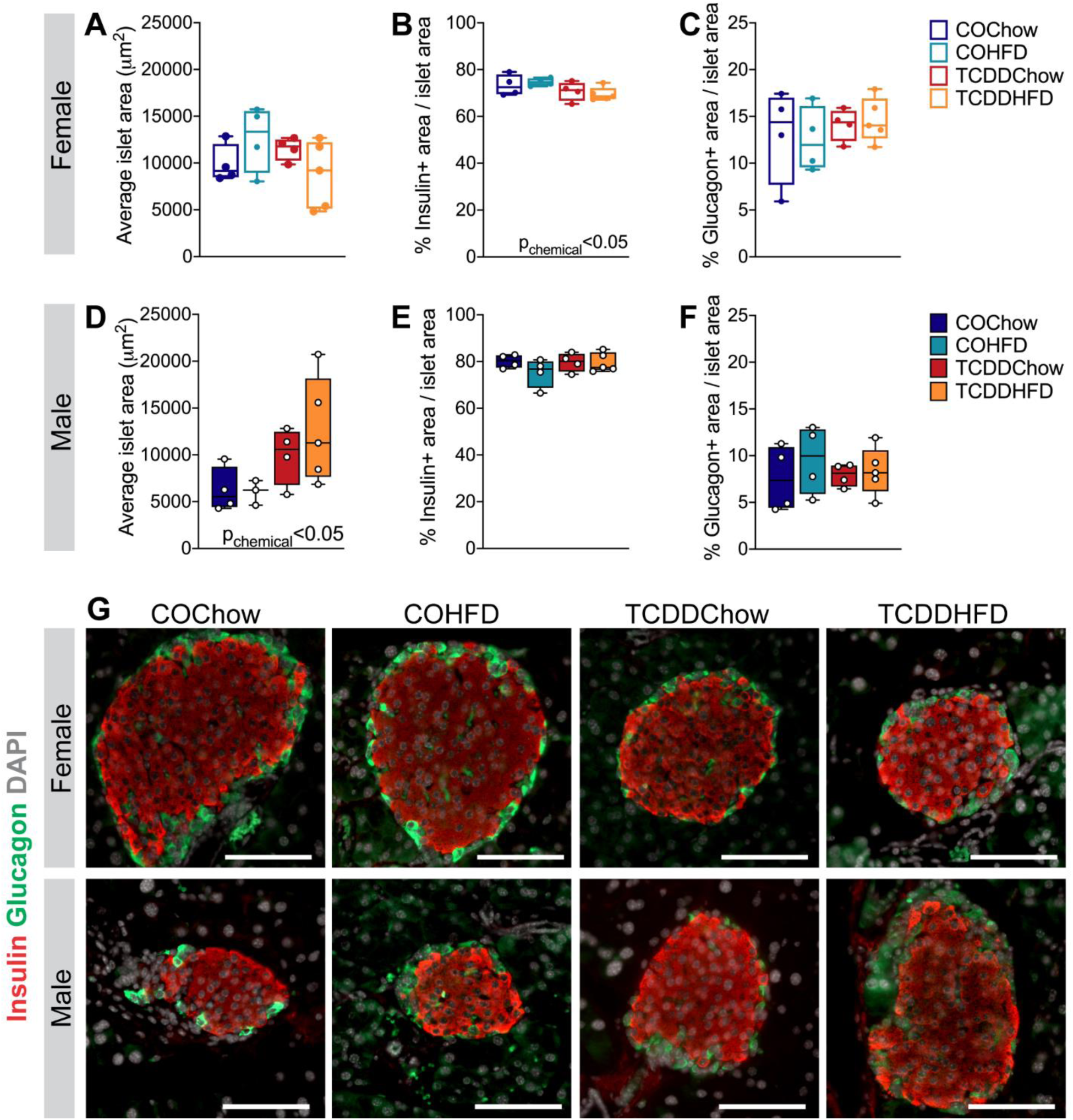
TCDD exposure has sex-specific effects on islet histology in mice. Whole pancreas was harvested at week 12 for analysis by immunofluorescence staining (see **Fig. 1A** for study timeline). (**A, D**) Average islet area, (**B, E**) % insulin^+^ area / islet area, (**C, F**) % glucagon^+^ area / islet area in (**A-C**) females and (**D-F**) males (n=4-5 per group). (**G**) Representative images of pancreas sections showing immunofluorescence staining for insulin and glucagon. Scale bar = 100 µm. All data are presented as box and whisker plots with median and min/max values. Individual data points on box and whisker plots represent biological replicates (different mice). The following statistical tests were used: two-way ANOVA with Tukey’s multiple comparison test.

### 3.5 TCDDHFD female mice have enriched endocrine-related pathways in islets relative to COChow females

We next performed TempO-Seq® analysis on isolated islets from a subset of females at week 12 to assess whole transcriptomic changes that might contribute to the *in vivo* phenotype observed in TCDDHFD females. We first compared the effect of TCDD exposure on females fed either chow or HFD (i.e. TCDDChow vs COChow and TCDDHFD vs COHFD). There were 595 unique DEGs in TCDDChow islets versus only 121 unique DEGs in TCDDHFD islets compared to their respective CO controls (**Supp. Fig. 3A**) (28). Pathway analysis comparing TCDDChow to COChow is presented in **Fig. 5Bi** and discussed below. Surprisingly, KEGG pathway analysis did not reveal any significant pathway alterations in TCDDHFD islets compared to COHFD islets. However, we did find that *Slc2a2* and *G6pc2*, two genes essential for proper beta cell function, were the most downregulated genes in TCDDHFD islets compared to COHFD islets (**Supp. Fig. 3B**) (28). These findings prompted us to assess our TempO-Seq® data using a different approach to better understand TCDD-induced changes in the islet transcriptome.

**Fig. 5.**
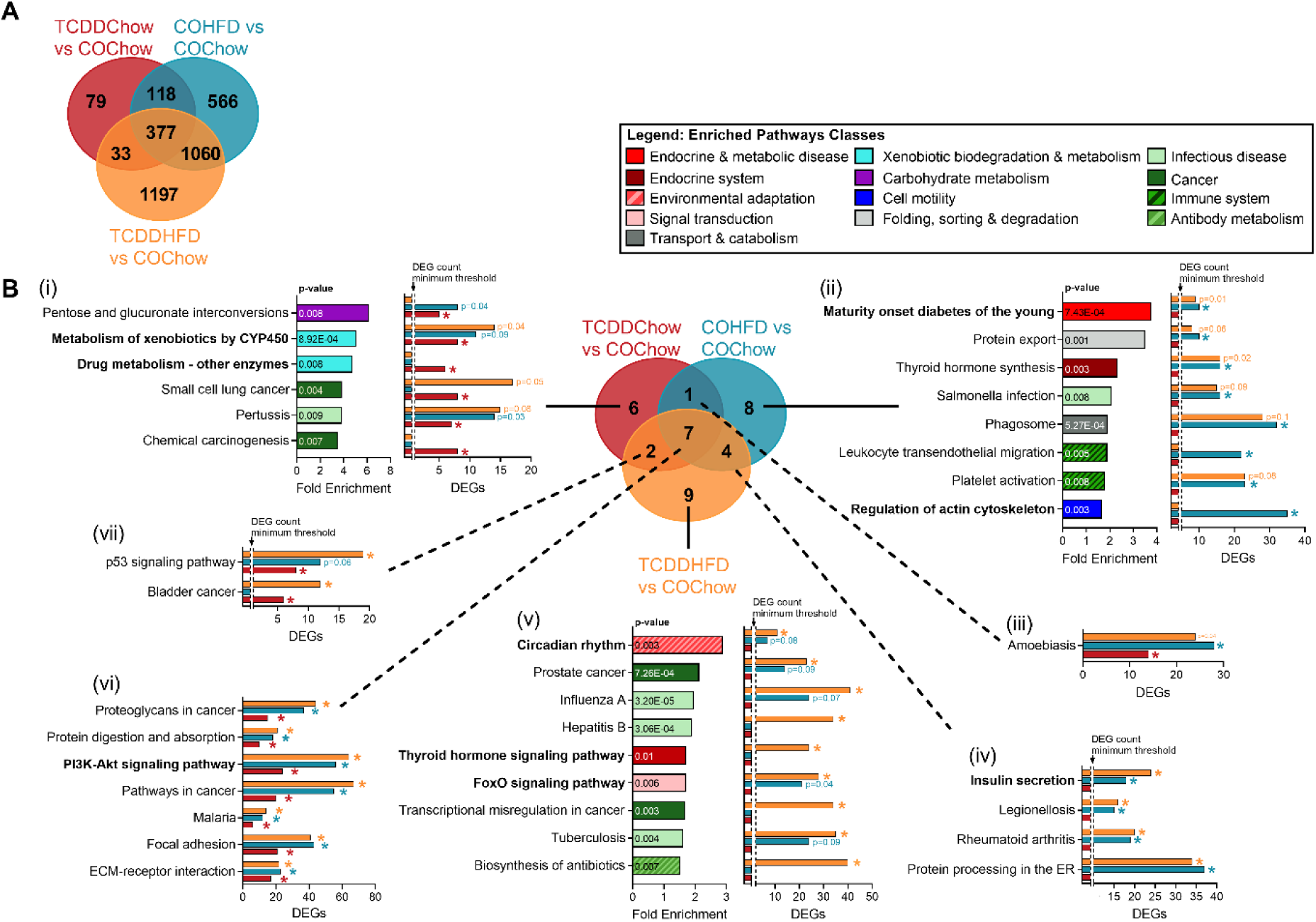
TCDDHFD female mice have enriched endocrine-related pathways relative to COChow females. Islets were isolated from females at week 12 of the study for TempO-Seq® analysis (n=3-4/group) (see **Fig. 1A** for study timeline). (**A**) Venn diagram displaying differentially expressed genes (DEGs) (adjusted p < 0.05, absolute fold change ≥ 1.5) in all experimental groups relative to COChow females. (**B**) KEGG pathway analysis was performed on all DEGs for each experimental group shown in (**A**). Results are displayed as a Venn diagram to show pathway overlap between experimental groups. For specific pathways, bar graphs show pathway fold enrichment in each group relative to COChow (DAVID modified Fisher Extract p-value < 0.01 relative to COChow females), and differentially expressed gene counts for statistically enriched pathways (coloured * = modified Fisher Extract p-value < 0.01 versus COChow). Pathways that were not trending or statistically enriched are indicated on bar graphs as not having the DEG count minimum threshold for significance.

We next compared all experimental groups to COChow islets to investigate the effect of either TCDD or HFD feeding alone (i.e. TCDDChow and COHFD, respectively, versus COChow) and in combination (TCDDHFD versus COChow). This allowed us to assess whether there is an interactive effect of HFD feeding and TCDD exposure on the islet transcriptome. TCDDChow islets had 607 DEGs, whereas COHFD islets had 2,121 DEGs and TCDDHFD islets had 2,667 DEGs compared to COChow islets (**Fig. 5A**). Interestingly, only 33 DEGs were unique to TCDD-exposed groups (i.e. TCDDChow and TCDDHFD) and unchanged in COHFD islets. In addition, only 79 genes were uniquely changed in TCDDChow islets (**Fig. 5A**). These results suggest that TCDD alone has relatively minor effects on gene expression in whole islets, which supports our *in vivo* observations (**Figs. 1-4**). In contrast, 1060 genes were altered in both HFD groups but not in the TCDDChow condition; 1,197 DEGs were unique to TCDDHFD islets and 566 DEGs unique to COHFD islets (**Fig. 5A**), suggesting that TCDD is driving abnormal changes in gene expression in response to HFD feeding.

We performed KEGG pathway analysis on all DEGs for each experimental group compared to COChow, as shown in **Fig. 5A** (i.e. TCDDChow-607 DEGs; COHFD-2,121 DEGs; TCDDHFD-2,667 DEGs). Since pathway enrichment is determined based on total number of DEGs and the number of DEGs varied between our experimental groups (**Fig. 5A**), we presented both pathway fold enrichment and number of DEGs within each pathway to avoid misinterpretation. Panel 5Bi shows pathways that were uniquely enriched in TCDDChow islets compared to COChow, thus representing the effects of TCDD alone on the islet transcriptome. As expected, “Xenobiotic Metabolism by CYP450” was highly enriched in TCDDChow islets compared to COChow. In fact, 50% of the pathways enriched in TCDDChow islets were involved in drug/chemical exposure, suggesting that TCDD alone mainly alters drug metabolism in islets. It is important to note that although “Xenobiotic Metabolism by CYP450” was not significantly enriched in COHFD or TCDDHFD islets compared to COChow, a greater number of genes in the pathway were significantly altered in HFD-fed than chow-fed females (**Fig. 5Bi**), suggesting that both TCDD exposure and HFD feeding alter this pathway.

We also assessed enriched pathways in COHFD and TCDDHFD islets compared to COChow controls to identify transcriptomic differences that may explain why TCDDHFD females lacked a compensatory increase in plasma insulin levels *in vivo*. “Maturity Onset Diabetes of the Young” (MODY) was the most enriched pathway in COHFD compared to COChow islets, but the number of DEGs from the MODY pathway was similar in both HFD groups irrespective of chemical treatment (**Fig. 5Bii**). In addition, “Insulin Secretion” was enriched in both HFD groups, although more genes were significantly different in TCDDHFD than COHFD compared to COChow islets (**Fig. 5Biv**). Interestingly, several pathways involved in beta cell function were differentially enriched in COHFD and TCDDHFD islets compared to COChow. First, “Regulation of Actin Cytoskeleton” was significantly altered in COHFD islets but not TCDDHFD islets, with 35 DEGs in the COHFD group (**Fig. 5Bii**). Therefore, changes in actin cytoskeleton in islets may be a normal response to HFD feeding that is absent in the TCDDHFD condition. Second, “Circadian Rhythm” was the most enriched pathway in TCDDHFD compared to COChow islets, and more genes within this pathway were changed in TCDDHFD than in COHFD islets (**Fig. 5Bv**). Likewise, “FoxO1 Signaling Pathway” and “Thyroid Hormone Signaling Pathway” were both only significantly enriched in TCDDHFD islets compared to COChow (**Fig. 5Bv**).

### 3.6 TCDDHFD females have inappropriate islet Cyp1a1 and circadian rhythm gene expression

We next used hierarchical clustering and heatmaps organized by experimental group (**Fig. 6, Supp. Fig. 4**) (28) to more carefully examine pathways of interest that were enriched in TCDDChow and TCDDHFD islets compared to COChow controls (**Fig. 5**). We were particularly interested in the “Xenobiotic Metabolism by CYP450” pathway since we have previously shown that TCDD induces CYP1A1 enzymes in islets (20,21). We expected to see a clear branching structure separating CO- and TCDD-exposed islets, but instead observed a main branch that separated COChow islets from all experimental groups, with overall gene expression being downregulated in all groups compared to COChow (**Fig. 6A, Supp. Fig. 4A**) (28). However, within the main branch, TCDDChow islets formed a sub-cluster, indicating slight differences in gene expression compared to HFD-fed females (**Fig. 6A**). Next, we specifically examined *Cyp1a1* expression as a marker of AhR activation by dioxin exposure (20,21) and found *Cyp1a1* was upregulated ~3.5-fold and ~3.2-fold in TCDDChow and COHFD islets, respectively, compared to COChow islets (**Fig. 6C**). In contrast, TCDDHFD islets showed a non-significant trend towards having only ~2-fold higher *Cyp1a1* compared to COChow islets, and *Cyp1a1* levels remained significantly lower than in TCDDChow islets (**Fig. 6C**). These results suggest that crosstalk between TCDD exposure and HFD feeding prevents normal *Cyp1a1* induction in islets; these findings were validated by qPCR (**Fig. 6D**). In addition, glutathione S-transferase (*Gstt1*), a phase II xenobiotic metabolism enzyme and another downstream target of AhR, was unchanged by HFD feeding or TCDD alone, but significantly downregulated in TCDDHFD islets compared to all other groups (**Fig. 6E**), further supporting an interaction between TCDD and HFD on xenobiotic metabolism pathways in islets.

**Fig. 6.**
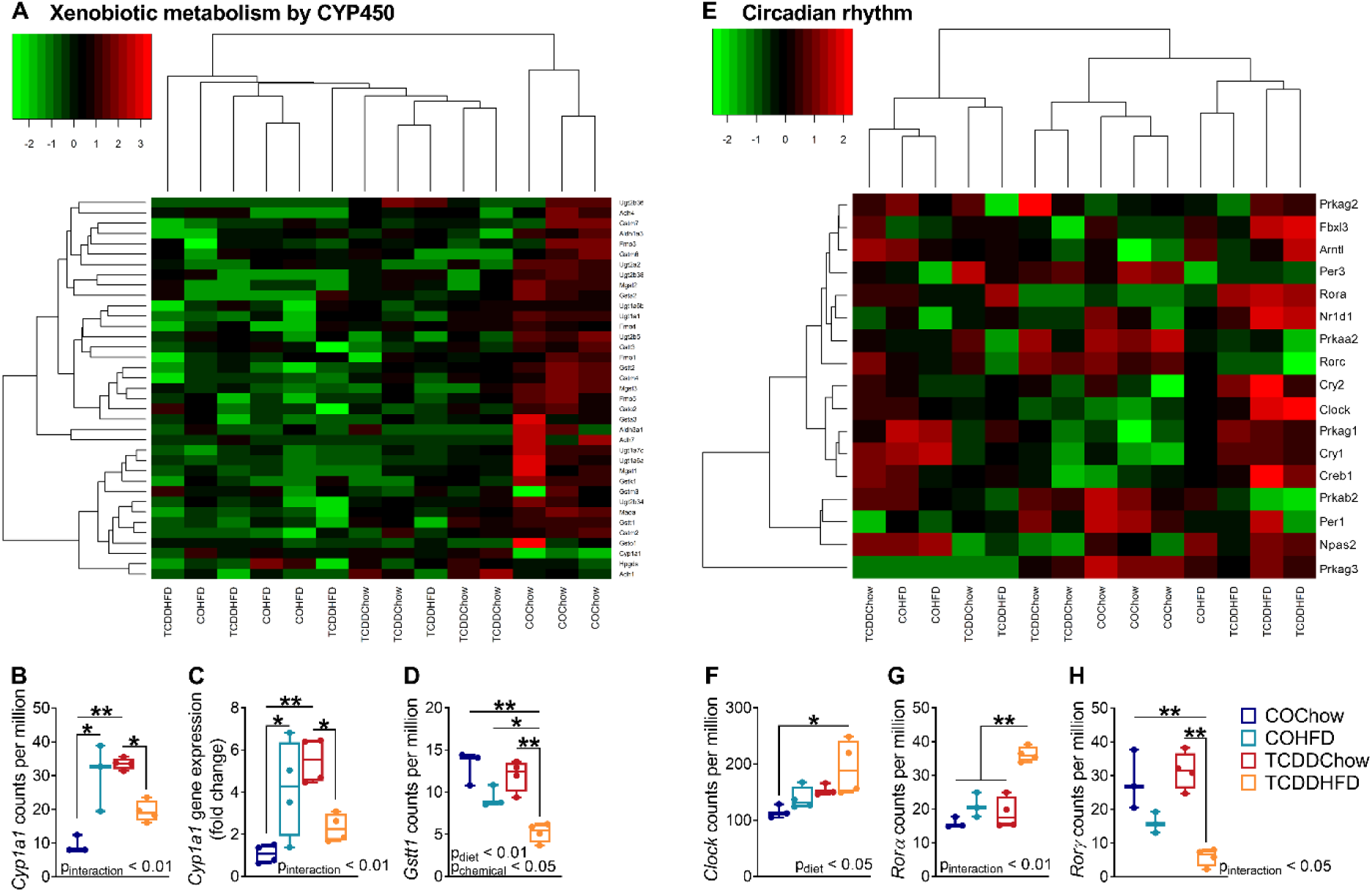
TCDDHFD females have inappropriate Cyp1a1 and circadian rhythm gene expression. Islets were isolated from females at week 12 of the study for TempO-Seq® and qPCR analysis (n=3-4/group) (see **Fig. 1A** for study timeline). (**A**) Hierarchal heatmap showing expression of genes involved in the “Xenobiotic Metabolism by CYP450” pathway. (**E**) Heatmap arranged by experimental group showing expression of genes involved in “Circadian Rhythm”. (**B, D, F-H**) Gene expression of (**B**) *Cyp1a1*, (**D**) *Gsst1*, (**F**) *Clock*, (**G**) *Rorα*, and (**H**) *Rorγ* in counts per million measured by TempO-Seq® analysis. (**C**) *Cyp1a1* gene expression measured by qPCR analysis. Individual data points on box and whisker plots represent biological replicates (different mice). All data are presented as median with min/max values. *p <0.05, **p <0.01. The following statistical tests were used: (**C-G**) two-way ANOVA with Tukey’s multiple comparison test.

We also assessed circadian rhythm gene expression patterns since this pathway was the most highly enriched in TCDDHFD islets (**Fig. 5v**). Unlike other pathways analysed in this study, the hierarchical heatmap for “Circadian Rhythm” was unclear (**Supp. Fig 4B**) (28). We observed two main branches in the hierarchical heatmap. The right branch clustered based on diet with HFD mice separated from chow mice, and showed further grouping within the diet subclusters that was driven by chemical exposure (**Supp. Fig 4B**) (28). However, the left branch of the hierarchal plot comprised all experimental groups and showed no clear pattern of clustering, preventing us from conclusively identifying overall changes in circadian rhythm (**Supp. Fig 4B**) (28). Instead, we used a heatmap organized by experimental group (**Fig. 6F**) and identified key regulators of circadian rhythm that were uniquely changed in TCDDHFD islets. For example, *Clock* was significantly upregulated in TCDDHFD islets compared to COChow islets (**Fig. 6E,F**). In addition, *Rorα* (retinoic acid receptor-related orphan receptor alpha), a member of the core clock machinery and a regulator of lipid/glucose homeostasis (34), was upregulated in TCDDHFD islets compared to all other experimental groups (**Fig. 6E,G**). Lastly, *Rorγ*, another regulator of the circadian clock (34), was downregulated in TCDDHFD females compared to chow-fed females (**Fig. 6E,H**). Taken together, these results suggest that TCDD exposure alters circadian rhythm in islets from HFD-fed mice.

### 3.7 TCDD exposure promotes HFD-induced changes in amino acid metabolism genes in islets

The *in vivo* results from the current study (**Figs. 2-3**) and previous findings from a separate study in pregnant mice (22) suggest that low-dose exposure to TCDD accelerates HFD-induced changes in metabolism in female mice. As such, we next compared the effects of HFD feeding on the islet transcriptome of females with either CO or TCDD background exposure (i.e. COHFD versus COChow and TCDDHFD versus TCDDChow). Interestingly, COHFD islets had 1,674 unique DEGs relative to COChow, whereas TCDDHFD islets had 706 unique DEGs relative to TCDDChow (**Fig. 7A**), indicating that the effect of HFD feeding varies substantially depending on chemical exposure. Indeed, KEGG pathway analysis revealed clear transcriptomic differences in islets from HFD-fed mice when on CO versus TCDD background (**Fig. 7B**). As shown in **Fig. 5B**, MODY was the most enriched pathway in COHFD versus COChow islets; this pathway was also altered in TCDDHFD versus TCDDChow islets, but to a lesser extent (**Fig. 7Bi**). “Regulation of Actin Cytoskeleton” and “PI3K-Akt Signaling Pathway” were only altered by HFD feeding when mice were exposed to CO (**Fig. 7Bi**), suggesting that TCDD exposure prevents these normal responses to HFD feeding. Most interestingly, the top two most enriched pathways in TCDDHFD islets compared to TCDDChow were “Alanine, Aspartate & Glutamate metabolism” and “Tyrosine Metabolism” (**Fig. 7Bii**). In fact, 40% of significantly enriched pathways in TCDDHFD islets are involved in amino acid metabolism (**Fig. 7Bii**), suggesting that HFD feeding causes abnormal changes to amino acid metabolism in islets from TCDD-exposed females.

**Fig. 7.**
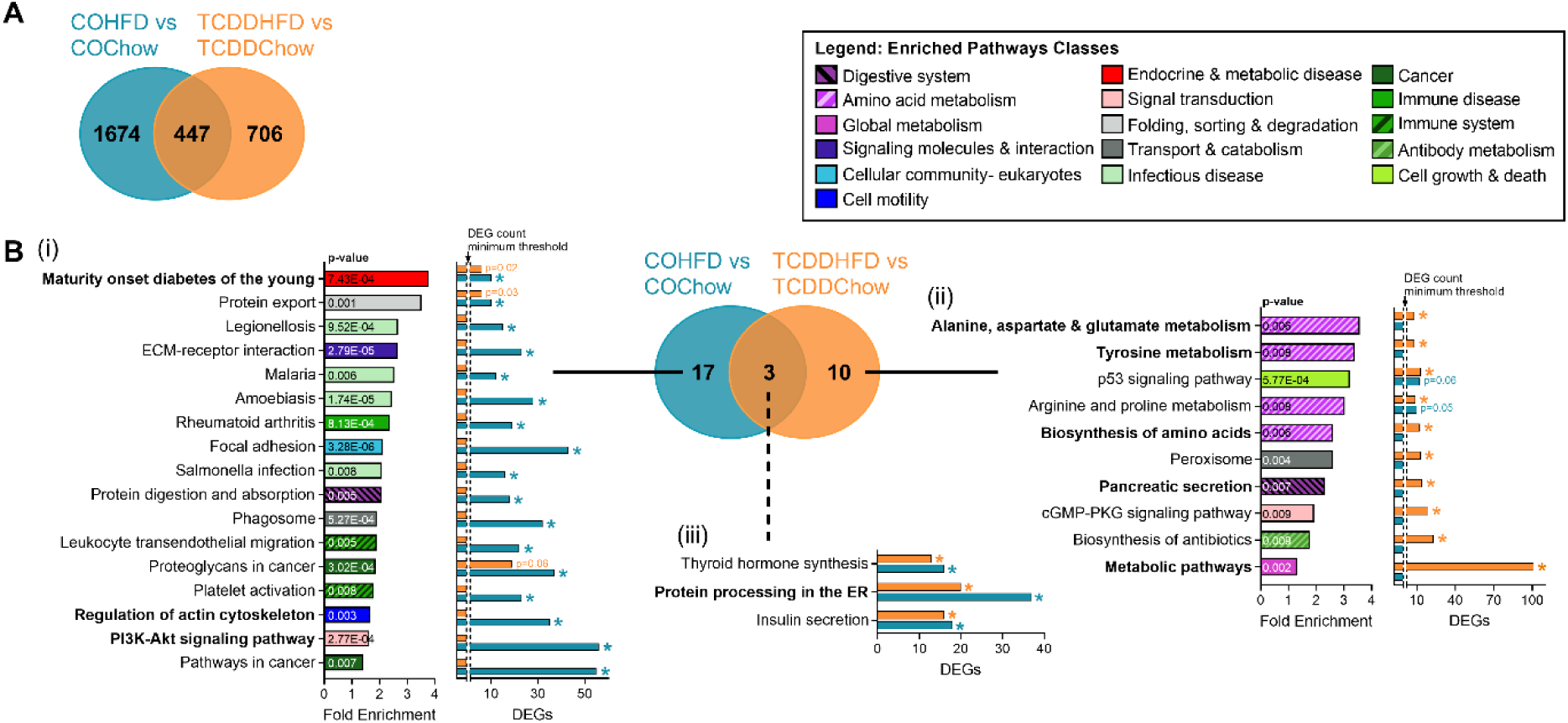
TCDD exposure promotes HFD-induced changes in amino acid metabolism genes in islets. Islets were isolated from females at week 12 of the study for TempO-Seq® analysis (n=3-4/group) (see **Fig. 1A** for study timeline). (**A**) Venn diagram displaying differentially expressed genes (DEGs) (adjusted p < 0.05, absolute fold change ≥ 1.5) in HFD fed females relative to their respective chow-fed control female. (**B**) KEGG pathway analysis was performed on all the DEGs for each comparison shown in (**A**). Results are displayed as a Venn diagram to show pathway overlap between experimental groups. For specific pathways, bar graph show pathway fold enrichment in each HFD-fed groups with respect to their chow-fed control (DAVID modified Fisher Extract p-value < 0.01 relative to respective chow-fed control), and differentially expressed gene counts for statistically enriched pathways (coloured * = modified Fisher Extract p-value < 0.01 versus chow-fed control). Pathways that were not trending or statistically enriched are indicated on bar graphs as not having the DEG count minimum threshold for significance.

### 3.8 HFD-induced changes in MODY and alanine, aspartate, and glutamate metabolism gene expression are altered by TCDD exposure

We generated heatmaps of the most enriched pathway in COHFD and TCDDHFD islets with respect to their chow-fed controls (pathway enrichment shown in **Fig. 7B**). Interestingly, the “MODY” heatmap showed that TCDDHFD islets form a unique cluster and generally had lower gene expression compared to all other groups (**Fig. 8A, Supp. Fig. 4C**) (28), suggesting an interactive effect of HFD feeding and TCDD exposure on expression of islet-specific genes. Furthermore, we noticed a distinct pattern in genes essential for maintaining beta cell function (**Fig. 8A-G, Supp. Fig. 4C**) (28). *MafA* and *Slc2a2* were significantly downregulated in TCDDHFD islets compared to COChow and TCDDChow islets, whereas expression only trended towards being downregulated in COHFD islets compared to COChow (**Fig. 8B,D**). *MafA* results were validated by qPCR (**Fig. 8C**). Similarly, *Hnf4a* was downregulated by HFD feeding, but this effect was worsened in TCDD-exposed islets (**Fig. 8E**), whereas *Pax6* was only downregulated in TCDDHFD islets compared to COChow (**Fig. 8F**). Interestingly, *Nkx6.1* was upregulated in COHFD compared to COChow islets, but downregulated in TCDDHFD islets compared to both COHFD and TCDDChow islets (**Fig. 8G**). These results confirm that TCDD exposure alters diet-induced changes in beta cell specific genes.

**Fig. 8.**
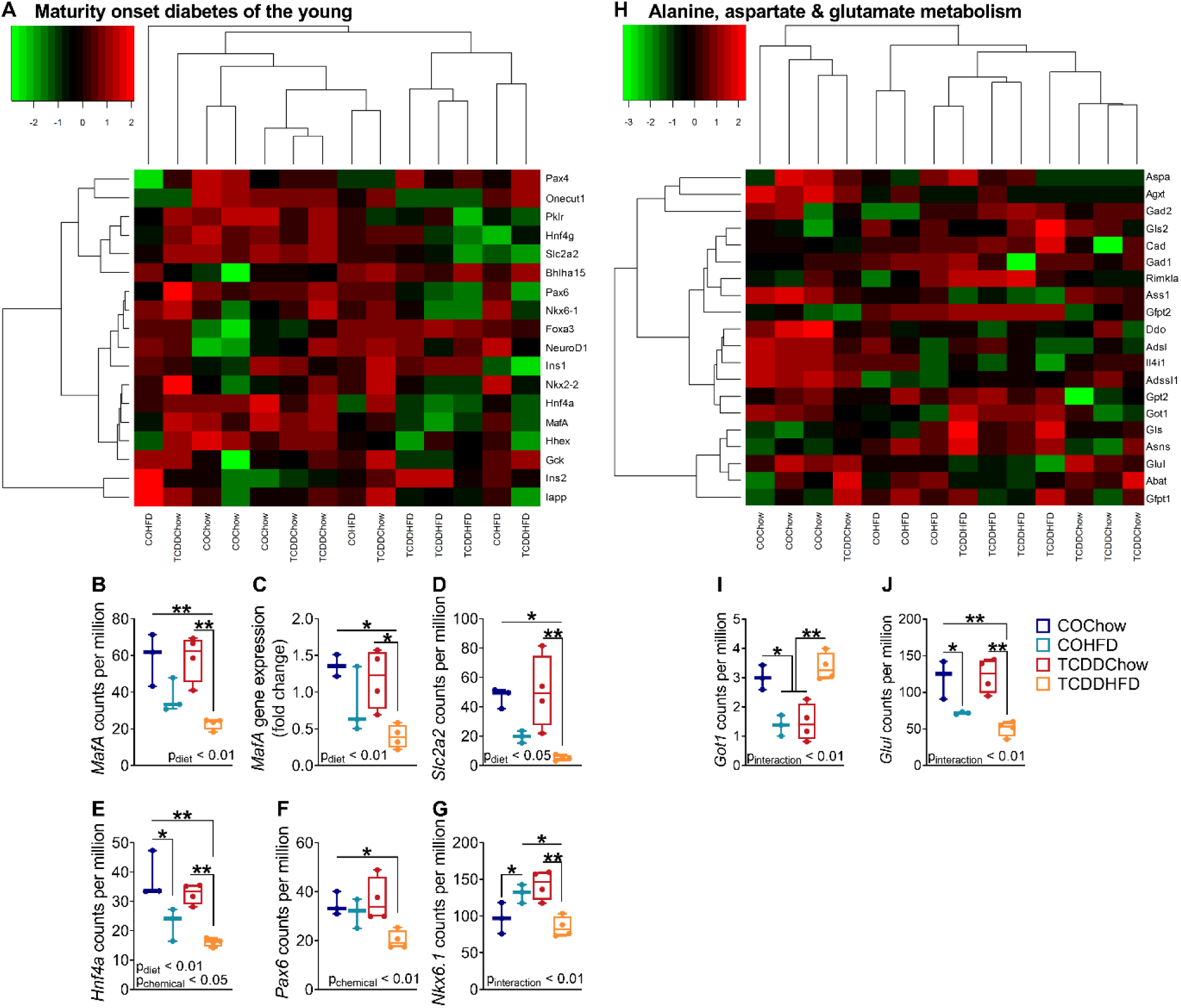
HFD-induced changes in MODY and alanine, aspartate, and glutamate metabolism gene expression are altered by TCDD exposure. Islets were isolated from females at week 12 of the study for TempO-Seq® and qPCR analysis (n=3-4/group) (see **Fig. 1A** for study timeline). (**A, H**) Hierarchical heatmaps showing expression of genes involved in (**A**) “Maturity Onset Diabetes of the Young” and (**H**) “Alanine, Aspartate and Glutamate Metabolism”. (**B, D-J**) Gene expression of (**B**) *MafA*, (**D**) *Slc2a2*, (**E**) *Hnf4α*, (**F**) *Pax6*, (**G**) *Nkx6.1*, (**I**) *Got1*, and (**J**) *Glul* in counts per million measured by TempO-Seq® analysis. (**C**) *MafA* gene expression measured by qPCR analysis. Individual data points on box and whisker plots represent biological replicates (different mice). All data are presented as median with min/max values. *p <0.05, **p <0.01. The following statistical tests were used: (**C-H**) two-way ANOVA with Tukey’s multiple comparison test.

The “Alanine, Aspartate and Glutamate Metabolism” pathway heatmap showed that COHFD and TCDD-exposed females formed a separate cluster compared to COChow females, suggesting that both HFD feeding and TCDD exposure alter amino acid metabolism (**Fig. 8H, Supp. Fig. 4D**) (28). Interestingly, TCDDHFD females formed a separate subcluster compared to both TCDDChow and COHFD females, suggesting that TCDD alters HFD-induced changes in amino acid metabolism (**Fig. 8H**). For example, aspartate aminotransferase (*Got1*) was significantly downregulated in COHFD and TCDDChow islets compared to COChow, but unchanged in TCDDHFD islets (**Fig. 8I**). In addition, glutamine synthase (*Glul*) was downregulated by HFD feeding alone, but this effect was worsened in TCDDHFD islets (p_interaction_ <0.01; **Fig. 8J**).

## 4. Discussion

Our study demonstrates that the effects of physiologically relevant low-dose TCDD exposure on glucose homeostasis are sex-dependent. In this 12-week study, low-dose TCDD alone did not impact glucose tolerance, insulin sensitivity, or plasma insulin levels in either chow-fed male or female mice, or in HFD-fed male mice. In contrast, HFD-fed females exposed to TCDD had accelerated onset of hyperglycemia and impaired compensatory hyperinsulinemia *in vivo* compared to HFD-fed females exposed to vehicle (CO). These data are in line with our previous findings that female mice exposed to low-dose TCDD transiently during pregnancy/lactation and subsequently fed HFD later in life had accelerated onset of hyperglycemia and suppression of glucose-induced plasma insulin levels compared to COHFD females (22). Islet morphometric analysis did not explain the phenotype observed in TCDDHFD females in the current study, but interesting interactions between HFD and TCDD exposure were seen at the transcriptomic level in isolated islets, including notable abnormal changes to endocrine and metabolic pathways. Although TCDD-exposed males did not display adverse metabolic outcomes, they had a modest increase in average islet size compared to CO-exposed mice, irrespective of diet. On the other hand, TCDD-exposed female mice rapidly developed HFD-induced hyperglycemia with no change to islet size, but instead showed reduced % beta cell area within islets. This differed from our previous findings with a single high dose (20 µg/kg) of TCDD, which reduced % beta cell area in males only and had no effect on islet size in either sex, suggesting that the effects of TCDD on islet morphology are dose-dependent. Regardless, overall effects of TCDD on glucose homeostasis were consistent between our two dosing protocols, with only female mice becoming glucose intolerant. These findings suggest that females are more susceptible to maladaptive metabolic responses following dioxin exposure. It is important to note that we did not conduct in-depth molecular analysis of male islets in this study so we cannot rule out the possibility that TCDD has deleterious molecular effects on male islets that might impair glucose homeostasis in a longer-term study.

Whether the TCDD-induced decrease in % beta cell area contributed to the inefficient adaptation of TCDD-exposed females to HFD feeding remains unclear. However, RNAseq analysis showed deficiencies in key beta cell genes in TCDDHFD islets compared to COChow that suggest defects at the beta cell level are contributing to this phenotype in female mice. For example, TCDD exposure altered HFD-induced changes in MODY genes such as *MafA, Hnf4α*, and *Slc2a2*, which all showed a more pronounced decrease in TCDDHFD islets than COHFD islets compared to chow-fed controls, and *Pax6*, which was downregulated only in TCDDHFD islets compared to COChow. These findings are consistent with our pregnancy model that showed a reduced proportion of beta cells expressing MAFA in the pancreas of TCDDHFD females compared to COHFD females (22). The beta cell transcription factors MAFA, PDX1, HNF4α, and PAX6 form an essential network for maintaining beta cell identity and regulating genes involved in insulin secretion (e.g. *Slc2a2*); inactivation of these genes is associated with beta cell dedifferentiation, metabolic inflexibility, and diabetes risk (35,36). Taken together, our results suggest that females exposed to prolonged low-dose TCDD have impaired metabolic adaptability, which may be linked to loss of beta cell identity, although single cell RNAseq analysis would be required to properly assess maturation status of beta cells. In addition, beta cell function was not directly measured in this study, but only glucose-induced plasma insulin levels were assessed. As such, it remains unclear whether these changes in gene expression are associated with reduced insulin secretion. Detailed *ex vivo* analysis of isolated islets is required to further elucidate the interaction between TCDD exposure and HFD feeding on beta cell function. Transcriptomic analysis at earlier timepoints are also required to understand acute effects of low-dose TCDD with/without HFD feeding on the islet phenotype.

Our TempO-Seq® analysis revealed “Metabolism of Xenobiotics by CYP450” as a highly enriched pathway in TCDD-exposed islets from female mice. This is consistent with our previous studies showing that single high-dose and multiple low-dose TCDD exposure induced CYP1A1 gene expression and enzyme activity in islets from male mice (20,21). The present study demonstrates for the first time that *Cyp1a1* is also upregulated in female islets following low-dose TCDD exposure. We also found that HFD feeding induces *Cyp1a1* expression in islets to a similar degree as TCDD exposure, suggesting a non-conventional role for CYP enzymes in islets. These findings are supported by our previous work showing that male *Cyp1a1/1a2* global knockout mice had suppressed glucose-stimulated insulin secretion in isolated islets *ex vivo* compared to wildtype controls (20). In the current study, the TCDD- and HFD-induced increase in *Cyp1a1* was suppressed in TCDDHFD females, suggesting cross-talk between TCDD and HFD feeding. This is similar to our finding that TCDD alone upregulated *CYP1A1* ~26-fold in human islets, but this induction was completely prevented by cotreatment with cytokines (20). Given that HFD feeding increases cytokine signalling (37), we speculate that there may be an interaction between AhR and cytokine signalling pathways in this mouse model. Whether impaired *Cyp1a1* induction contributes to the metabolic phenotype observed in TCDDHFD females remains unclear.

TempO-Seq® analysis revealed additional unique transcriptomic changes in metabolic and endocrine pathways in TCDDHFD islets compared to COChow or TCDDChow islets that further support a TCDD-driven defect in metabolic adaptability. We focus on two key pathways of interest here, although future studies should further investigate the interactive effect of TCDD and HFD on other pathways that were enriched in our study, including “Thyroid Signalling” and “Foxo1 Signaling”. Our analysis revealed changes in the “Circadian Rhythm” pathway in islets from TCDDHFD females, including alterations in key islet circadian rhythm. These data are interesting given that alterations to the circadian clock have been associated with diabetes, obesity, and metabolic syndrome in humans (38–40). Abnormal expression of clock genes has also been reported in islets from type 2 diabetic donors compared to non-diabetic donors, and insulin content was positively correlated with expression of clock genes, indicating that disruptions to the islet circadian clock contributes to beta cell dysfunction and diabetes pathogenesis (41,42). In fact, knockdown of *Clock* in human islets led to dysfunctional glucose-stimulated insulin secretion and altered expression of genes involved in insulin secretion (43). Islet-specific *Clock* knockout mice also exhibit hypoinsulinemia, hyperglycemia, and an impaired insulin secretory response (44). Lastly, experimental disruption of circadian rhythms accelerates the onset of hyperglycemia and loss of functional beta cells in diabetes-prone rats (45). Therefore, it is possible that the glucose intolerance and lack of compensatory hyperinsulinemia in TCDDHFD female mice involves altered islet circadian rhythms, but more detailed analysis is required to understand the mechanism(s) involved.

TCDD exposure has previously been associated with changes to circulating amino acid concentrations (46–48) and hepatic amino acid metabolism genes *in vivo* (49) and *in vitro* (50,51). Our transcriptomic data indicate that TCDD may also alter amino acid metabolism in islets under conditions of HFD feeding, which could have important implications for both alpha cell and beta cell function. Amino acids are key regulators of glucagon secretion (52–55) and multiple studies have shown that alanine and glutamine, specifically, are potent glucagon secretagogues (56–58). Amino acids also enhance glucose-stimulated insulin secretion (58–61) and promote the expression of genes involved in beta cell signal transduction, metabolism, and insulin secretion (60,61). Perturbations to circulating amino acid levels have been associated with type 2 diabetes, obesity, and insulin resistance (62,63). Therefore, the finding that 40% of enriched pathways in HFD-fed females are involved in amino acid metabolism, but only when combined with background TCDD exposure, likely has important implications for islet function and overall glucose homeostasis. Further investigation is required to assess the interactive effect of TCDD and HFD on both glucagon and insulin secretion.

To our knowledge this is the first study to do a head-to-head comparison of the effects of low-dose TCDD exposure on diabetes risk in male and female mice. It is important to note that mice were exposed to TCDD through i.p. injection rather than the more physiological route of oral exposure for logistical reasons. A caveat of this approach is bypassing the effects of TCDD on the gut and gut hormones (including GLP-1 and GIP), which are important modulators of insulin secretion. As such, this may have diminished the impact of TCDD on glucose homeostasis, however further research is required to assess whether TCDD alters incretin hormones. Despite the route of exposure, our findings are consistent with epidemiological evidence showing that females with high serum pollutant levels have a higher risk of developing diabetes than males (5,10,17,18). Our mouse study shows that 12 weeks of low-dose TCDD alone does not cause overt diabetes but does increase susceptibility to diet-induced hyperglycemia in female mice, emphasizing the need to study the interaction of pollutants with other metabolic stressors such as diet and the importance of considering sex differences in studies about pollutant-induced diabetes risk.

## Abbreviations

AhR: Aryl hydrocarbon receptor
CO: Corn oil
Cyp1a1: Cytochrome P450 1A1
DEG: Differentially expressed gene
GTT: Glucose tolerance test
HFD: High-fat diet
ITT: Insulin tolerance test
POPs: Persistent organic pollutants
TCDD: 2,3,7,8-tetrachlorodibenzo-*p*-dioxin

## Acknowledgements

We thank Salar Farokhi Boroujeni and Andrea Rowan-Carroll for their technical assistance with sample analysis for this project. This research was supported by a Canadian Institutes of Health Research (CIHR) Project Grant (#PJT-2018-159590), Natural Sciences and Engineering Research Council of Canada (NSERC) Discovery Grant (RGPIN-2017-06272), the Canada Foundation for Innovation John R. Evans Leaders Fund (#37231), and an Ontario Research Fund award. M.P.H. was supported by an Ontario Graduate Scholarship (OGS) and CIHR CGS-D award. H.B. received a NSERC Undergraduate Student Research Award (USRA) and a Carleton University Walker Summer Research Award. J.E.B. is the guarantor for this work.

## Author Contributions

J.E.B. and G.M. conceived the experimental design. J.E.B., G.M., and M.P.H. wrote the manuscript. G.M., M.P.H., H.L.B., J.Z., S.F.B, K.R.C.R., A.W., R.G., J.K.B., C.L.Y., and J.E.B. were involved with acquisition, analysis, and interpretation of data. All authors contributed to manuscript revisions and approved the final version of the article.

