## Supplemental Data for "Prolonged low-dose dioxin exposure impairs metabolic adaptability to high-fat diet feeding in female but not male mice"

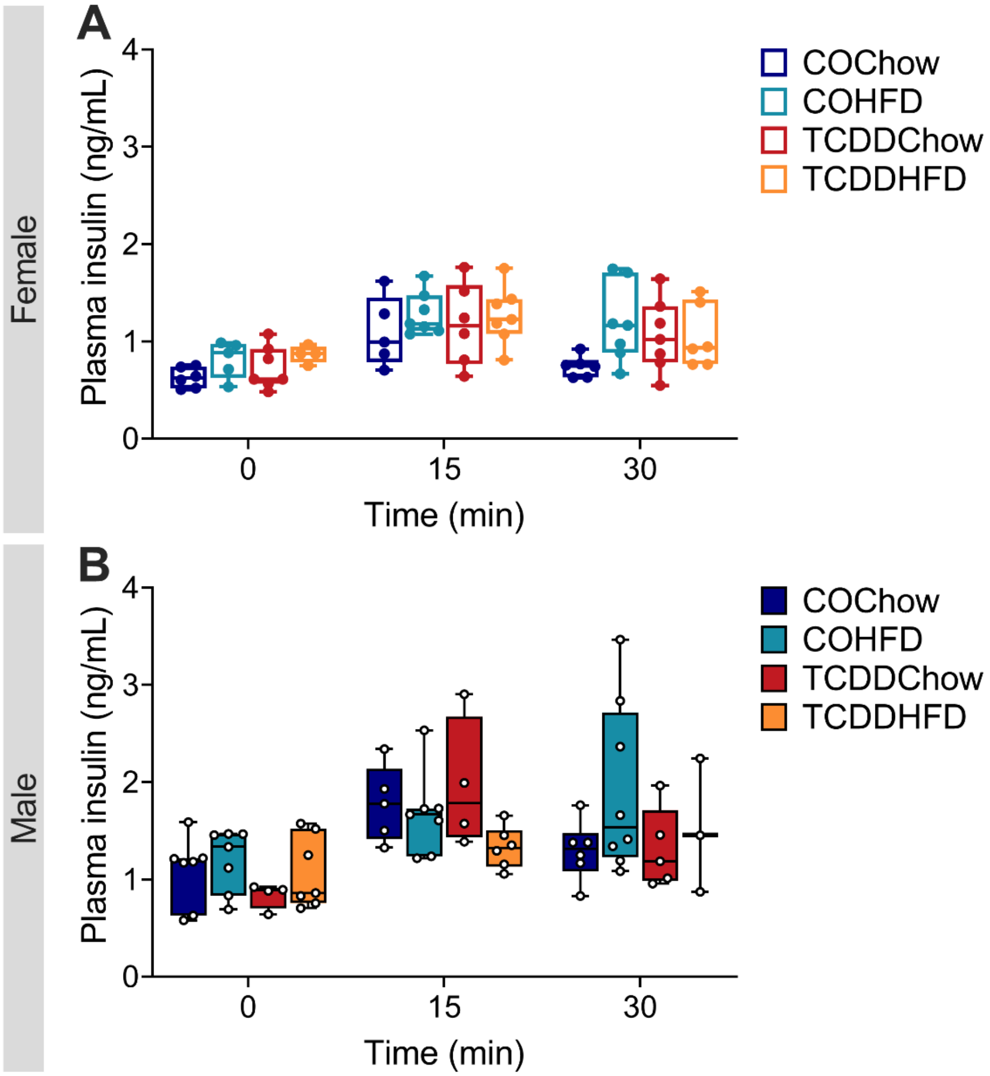


**Supp. Fig. 1 TCDD exposure did not alter glucose-induced plasma insulin levels at week 4.** Glucose-stimulated plasma insulin levels were assessed *in vivo* at week 4 in (**A**) females and (**B**) males (n=3-8 per group) (see **Fig. 1A** for study timeline). Data are presented as box and whisker plots. Individual data points on box and whisker plots represent biological replicates (different mice). The following statistical tests were used: (**A-B**) one-way ANOVA with Tukey's multiple comparison test at each time point.

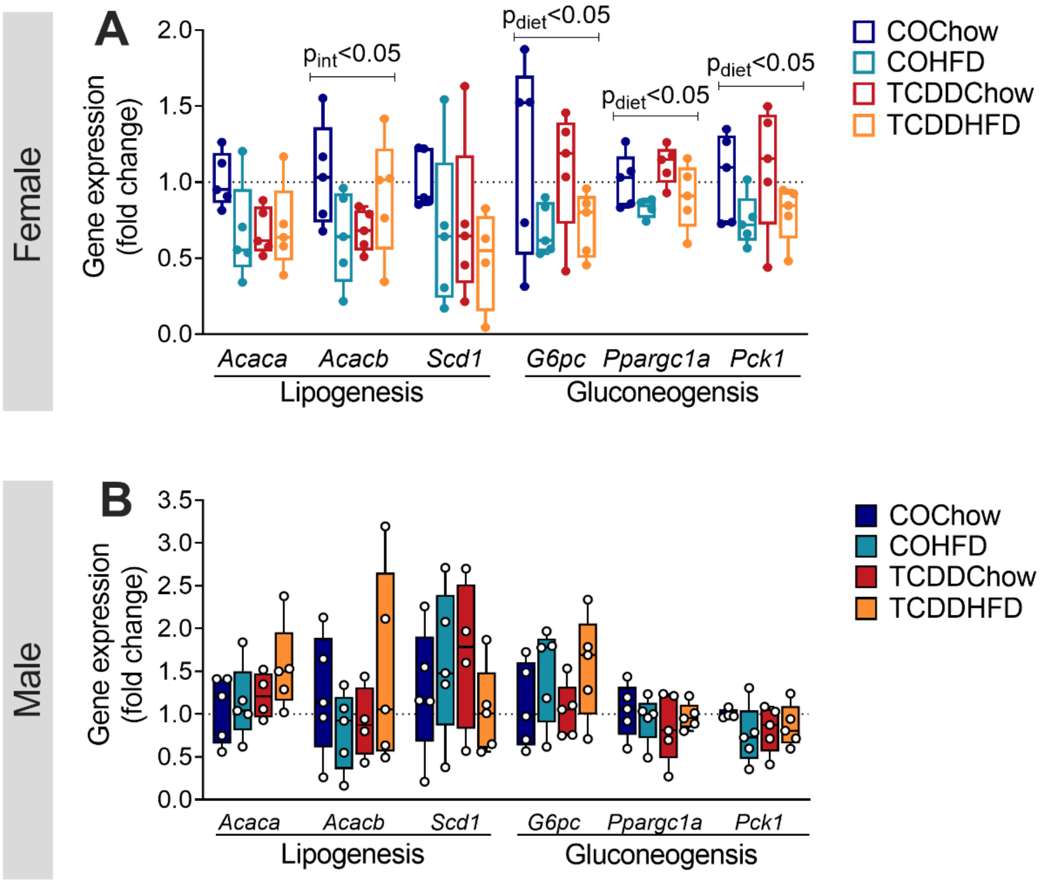


**Supp. Fig. 2 TCDD exposure did not alter hepatic expression of genes involved in insulin-dependent lipogenesis and gluconeogenesis**. Liver was harvested at week 12 of the study for gene expression analysis by qPCR (see **Fig. 1A** for study timeline). Expression of genes involved in lipogenesis and gluconeogenesis were measured in (**A**) females and (**B**) males. All data are presented as median with min/max values in box and whisker plots. Individual data points on box and whisker plots represent biological replicates (different mice). The following statistical tests were used: (**A-B**) two-way ANOVA with Tukey's multiple comparison test.

**A**

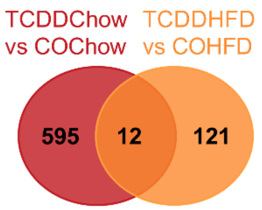


**B** Most downregulated genes

| Gene | Fold Change | p-value |
|---|---|---|
| *Slc2a2* | -3.69 | 1.29E-09 |
| *G6pc2* | -3.20 | 6.15E-15 |
| *Mrln* | -3.11 | 8.66E-05 |
| *Rbm3* | -2.85 | 2.70E-14 |
| *Igfbp3* | -2.68 | 6.22E-12 |

Most upregulated genes

| Gene | Fold Change | p-value |
|---|---|---|
| *Akr1b3* | 3.91 | 3.25E-21 |
| *Vmn1r121* | 2.67 | 0.001033 |
| *Tfdp2* | 2.46 | 2.19E-05 |
| *Gm648* | 2.41 | 0.000147 |
| *Ephx1* | 2.39 | 3.26E-06 |

**Supp. Fig. 3 TCDD-induced changes in gene expression relative to CO-exposed controls.** Female mice were injected with corn oil (CO) or 20 ng/kg/d TCDD 2x/week for 12 weeks, and either fed standard chow or 45% HFD (see **Fig. 1A** for study timeline). Islets were isolated at 12 weeks for TempO-Seq® analysis. (**A**) Venn diagram displaying differentially expressed genes (DEGs) (adjusted $p < 0.05$, absolute fold change ≥ 1.5) in TCDD-exposed females relative their respective CO-exposed control female. (**B**) Five most down- and up-regulated genes in TCDDHFD females when compared to COHFD females.

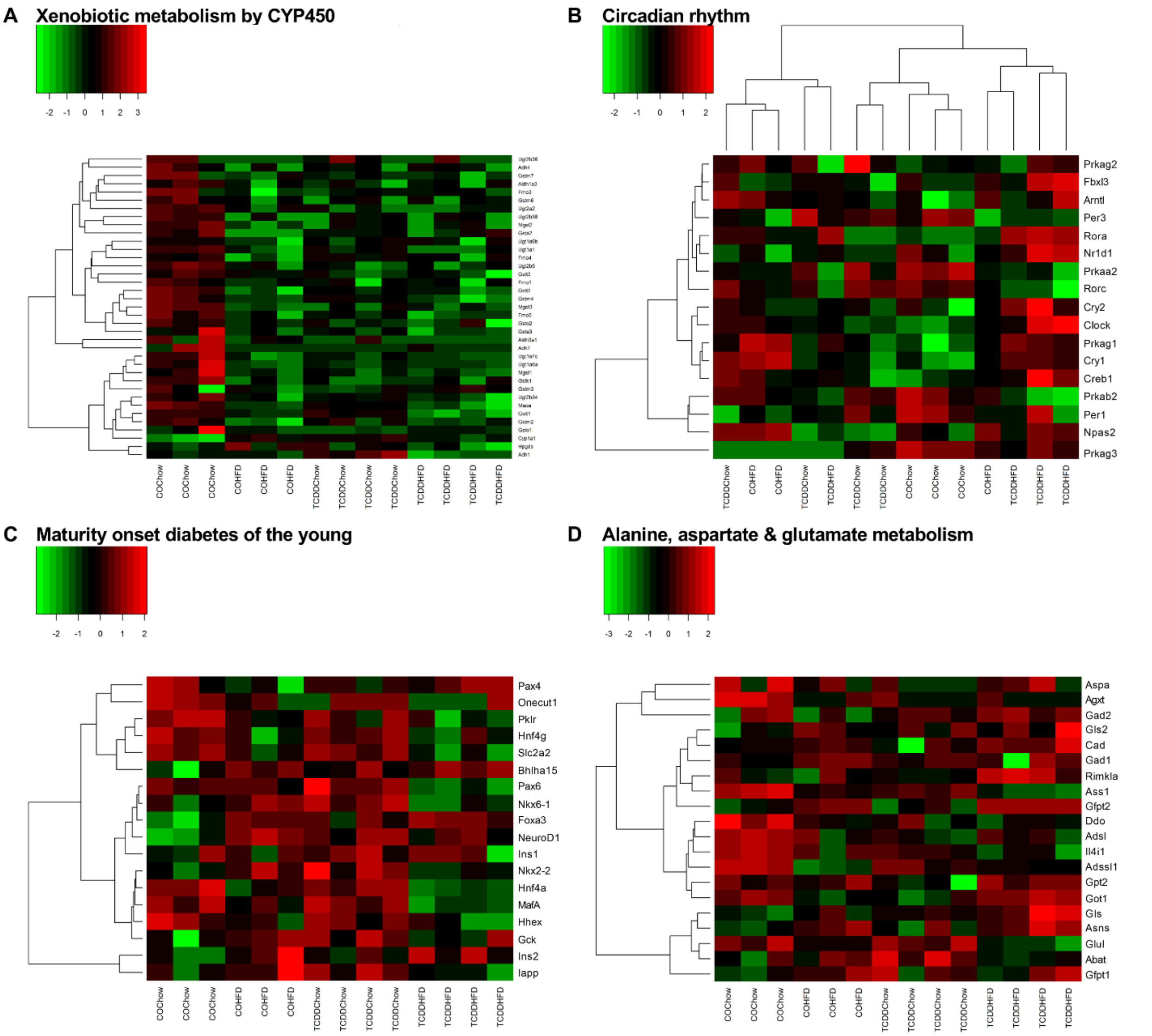


**Supp. Fig. 4 Heatmaps displaying gene expression profiles for "Xenobiotic Metabolism by CYP450", "Circadian Rhythm", "Maturity Onset Diabetes of the Young", and "Alanine, Aspartate & Glutamate Metabolism".** Female mice were injected with corn oil (CO) or 20 ng/kg/d TCDD 2x/week for 12 weeks, and either fed standard chow or 45% HFD (see **Fig. 1A** for study timeline). Islets were isolated at 12 weeks for TempO-Seq® analysis. Heatmaps arranged by experimental group showing expression levels of genes involved in (**A**) "Xenobiotic Metabolism by CYP450", (**C**) "Maturity Onset Diabetes of the Young", and (**D**) "Alanine, Aspartate and Glutamate metabolism". (**B**) Hierarchical heatmap showing "Circadian Rhythm" gene expression profile.

**Supp. Table 1:** qPCR primer sequences.

| **Target** | **Forward Primer** | **Reverse Primer** |
|---|---|---|
| *Acaca* | CCT GAC AAA CGA GTC TGG CT | CAT TCC ATG CAG TGG TCC CT |
| *Acacb* | TCC GTG CCT TTG TAC AGT CC | CAG GCT CCA AGT GGC GAT AA |
| *Cyp1a1* | ATC ACA GAC AGC CTC ATT GAG C | AGA TAG CAG TTG TGA CTG TGT C |
| *G6pc* | TAC TAC AGC AAC AGC TCC GTG | TCC CAA CCA CAA GAT GAC GTT |
| *Mafa* | AGT CGT GCC GCT TCA AG | CGC CAA CTT CTC GTA TTT CTC C |
| *Pck1* | TGG GAA CTC ACT ACT CGG GA | TTC TTC TTG CCT TCG GGG TT |
| *Ppargc1a* | CTT GAC TGG CGT CAT TCG G | CGC TAC ACC ACT TCA ATC CAC |
| *Ppia* | AGC TCT GAG CAC TGG AGA GA | GCC AGG ACC TGT ATG CTT TA |
| *Scd1* | CTG AAC ACC CAT CCC GAG AG | GTG GTG GTG GTC GTG TAA GA |
